# Metagenomic analysis of antimicrobial resistance, virulence, and mobile genetic elements in the gut microbiota of Caprinae species

**DOI:** 10.1101/2025.07.09.663846

**Authors:** Jin-Wen Su, Hany M. Elsheikha, Li Guo, Rui Liu, Kai-Meng Shang, Hai-Long Yu, He Ma, Hong-Bo Ni, Bei-Ni Chen, Xiao-Xuan Zhang, Xing Yang

**Affiliations:** Department of Medical Microbiology and Immunology, School of Basic Medicine, Dali University, Dali, Yunnan Province, PR China; College of Life Sciences, Changchun Sci-Tech University, Shuangyang, Jilin Province, PR China; College of Veterinary Medicine, Qingdao Agricultural University, Qingdao, Shandong Province, PR China; Faculty of Medicine and Health Sciences, School of Veterinary Medicine and Science, University of Nottingham, Sutton Bonington Campus, Loughborough, United Kingdom; Animal Science and Technology College, Jilin Agricultural Science and Technology University, Jilin, Jilin Province, PR China

**Keywords:** Caprinae species, Gut microbiome, Antimicrobial resistance genes, Virulence factor genes, Mobile genetic elements, Metagenomics

## Abstract

The gut microbiota of livestock serves as a reservoir for antimicrobial resistance (AMR), yet Caprinae species remain understudied in this context. In this comprehensive metagenomic study, we analyzed 779 gut samples from Caprinae animals and reconstructed 17,023 high-quality metagenome-assembled genomes (MAGs). From these, we identified 4,685 antimicrobial resistance genes (ARGs) and 5,401 virulence factor genes (VFGs). *Escherichia coli* emerged as a major host carrying high burdens of both ARGs and VFGs. Strong positive correlations between ARGs, VFGs, and mobile genetic elements (MGEs) suggest potential co-selection and genetic linkage. Although MGEs were found in only 1.45% of MAGs, 23 ARGs were physically co-located with MGEs, indicating mobility potential. Additionally, three ARGs were embedded within viral genomes, two of which were associated with *Myoviridae* phages and one with an unclassified viral source, implicating phages in AMR dissemination. Comparative analyses revealed 292 ARG types shared between Caprinae and the human gut microbiota, including 20 genes representing six clinically critical resistance types: *tetX1*, *tetX4*, *tmexD3*, *vanD*, *vanR*, and *vanS*—conferring resistance to tigecycline, vancomycin, and polymyxins. These findings expand our understanding of the resistome and virulome in Caprinae animals and highlight potential zoonotic transmission pathways, underscoring the need for targeted AMR surveillance and mitigation strategies.

## Introduction

Antimicrobial resistance (AMR) is an escalating global health crisis with profound implications for human medicine, animal husbandry, and ecological stability ^1–3^. Among the various contributors to this crisis, the gut microbiota of animals has emerged as a critical reservoir and conduit for AMR genes (ARGs), virulence factor genes (VFGs), and mobile genetic elements (MGEs). These microbial communities facilitate the emergence, enrichment, and horizontal transfer of resistance determinants within and between species, as well as into the environment ^4–6^. Understanding the distribution and transmission pathways of these genetic elements is therefore essential for assessing the risk of AMR spread across ecological and host boundaries.

Caprinae species—including domestic sheep (*Ovis aries*), goats (*Capra hircus*), and bharals (*Pseudois nayaur*)—occupy a broad range of ecological niches and play a vital role in global agriculture and rural livelihoods. Their diverse husbandry practices, close interaction with human populations, and ecological mobility position them as potentially significant players in the resistome landscape ^7^. While previous studies have characterized AMR in the gut microbiota of ruminants ^8,9^, comprehensive metagenomic investigations focused specifically on Caprinae species remain limited.

In addition to bacterial populations, bacteriophages and other gut-associated viruses may also influence AMR dynamics. Although phage-mediated transfer of ARGs is not yet considered a primary mechanism, growing evidence suggests that under certain ecological conditions, viruses can serve as ARG carriers and facilitate their dissemination ^10–14^. The presence of resistance genes in viral genomes within the gut ecosystem therefore warrants closer scrutiny.

In this study, we performed a large-scale metagenomic analysis of 779 gut samples from Caprinae animals to systematically characterize the distribution of ARGs, VFGs, and MGEs. We also assessed the presence of ARGs within the viral fraction of the microbiome. By integrating taxonomic and functional profiling, this work enhances our understanding of the role of Caprinae gut microbiota in shaping AMR ecology. Our findings provide a foundational framework for future investigations into resistance transmission and offer critical insights into the broader One Health implications of AMR dissemination in livestock-associated environments.

## Results

### A comprehensive MAG catalog of the caprinae gut microbiota

From a total of 779 gut metagenomic samples derived from Caprinae animals, we reconstructed 63,126 metagenome-assembled genomes (MAGs). Following stringent quality filtering (completeness ≥50%, contamination ≤10%) and dereplication at 99% average nucleotide identity (ANI), we curated a high-confidence dataset of 17,023 non-redundant MAGs (Fig. 1A; Supplementary Table 2). These MAGs demonstrated robust genomic quality, with completeness values ranging from 50% to 100% (mean: 76.95%) and contamination levels between 0% and 10% (mean: 2.09%) (Fig. 1B). The GC content spanned from 22.47% to 73.02% (mean: 47.27%), while genome sizes varied from 0.22 Mb to 8.15 Mb (mean: 1.92 Mb) (Fig. 1C), reflecting the phylogenetic and functional diversity within the dataset.

**Fig. 1:**
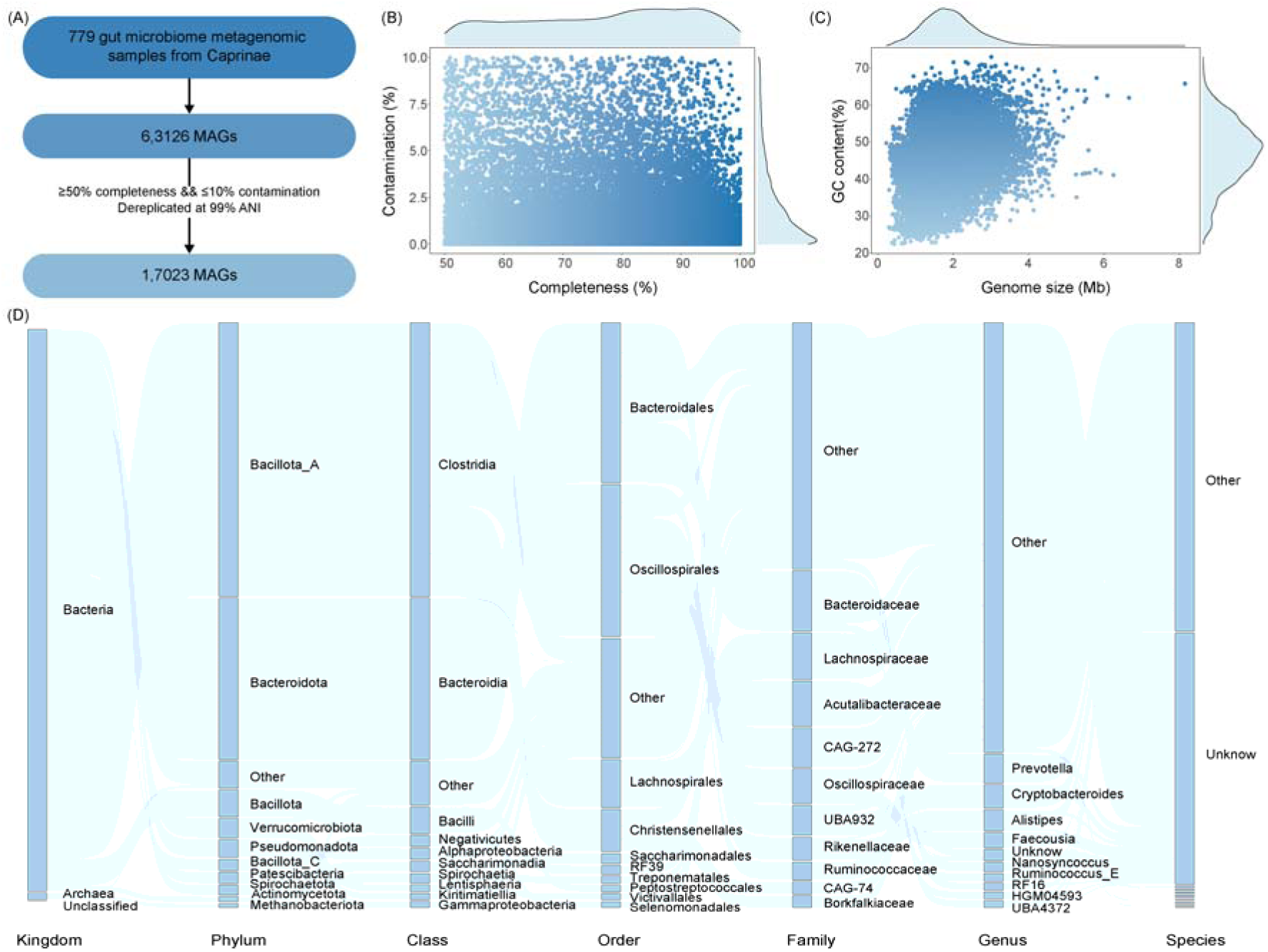
Comprehensive genomic landscape of the Caprinae gut microbiota. (A) Workflow illustrating the selection and quality filtering process of high-quality MAGs from Caprinae gut samples. (B) Scatter plot depicting completeness and contamination metrics for the 17,023 high-quality MAGs, with each dot representing a single MAG. (C) Distribution of GC content versus genome size across the MAG dataset. (D) Sankey diagram showing the taxonomic composition of the 17,023 MAGs across multiple taxonomic levels.

Taxonomic classification revealed one unclassified MAG, 218 archaeal MAGs, and the remainder of bacterial origin (Fig. 1D; Supplementary Table 2). These genomes collectively spanned 32 phyla, 45 classes, 109 orders, 260 families, 1,253 genera, and 3,612 species. Remarkably, 7,508 MAGs (44.11%) could not be assigned to any known species, underscoring a rich reservoir of previously uncharacterized microbial diversity in the Caprinae gut. At higher taxonomic levels, the phyla *Bacillota_A* (48.09%) and *Bacteroidota* (28.29%) were predominant. At the genus level, *Prevotella* (5.13%), *Cryptobacteroides* (4.10%), and *Alistipes* (3.67%) were most abundant, highlighting the dominance of anaerobic commensals commonly associated with herbivorous gut ecosystems.

### ARG landscape in the caprinae gut microbiota

To assess the prevalence and distribution of ARGs in the Caprinae gut microbiome, all 17,023 high-quality MAGs were screened against the Comprehensive Antibiotic Resistance Database (CARD). This analysis identified a total of 4,685 ARGs, with 2,709 MAGs (15.91%) harboring at least one ARG, spanning 337 unique resistance phenotypes (Supplementary Table 3). The most frequently detected resistance phenotypes were against tetracyclines (35.35%), elfamycins (29.13%), and multi-drug classes (10.98%) (Fig. 2A). The dominant resistance mechanisms included antibiotic target alteration (43.92%) and target protection (36.36%), followed by antibiotic inactivation (12.14%), efflux (3.54%), and multi-mechanism resistance (3.61%) (Fig. 2B).

**Fig. 2:**
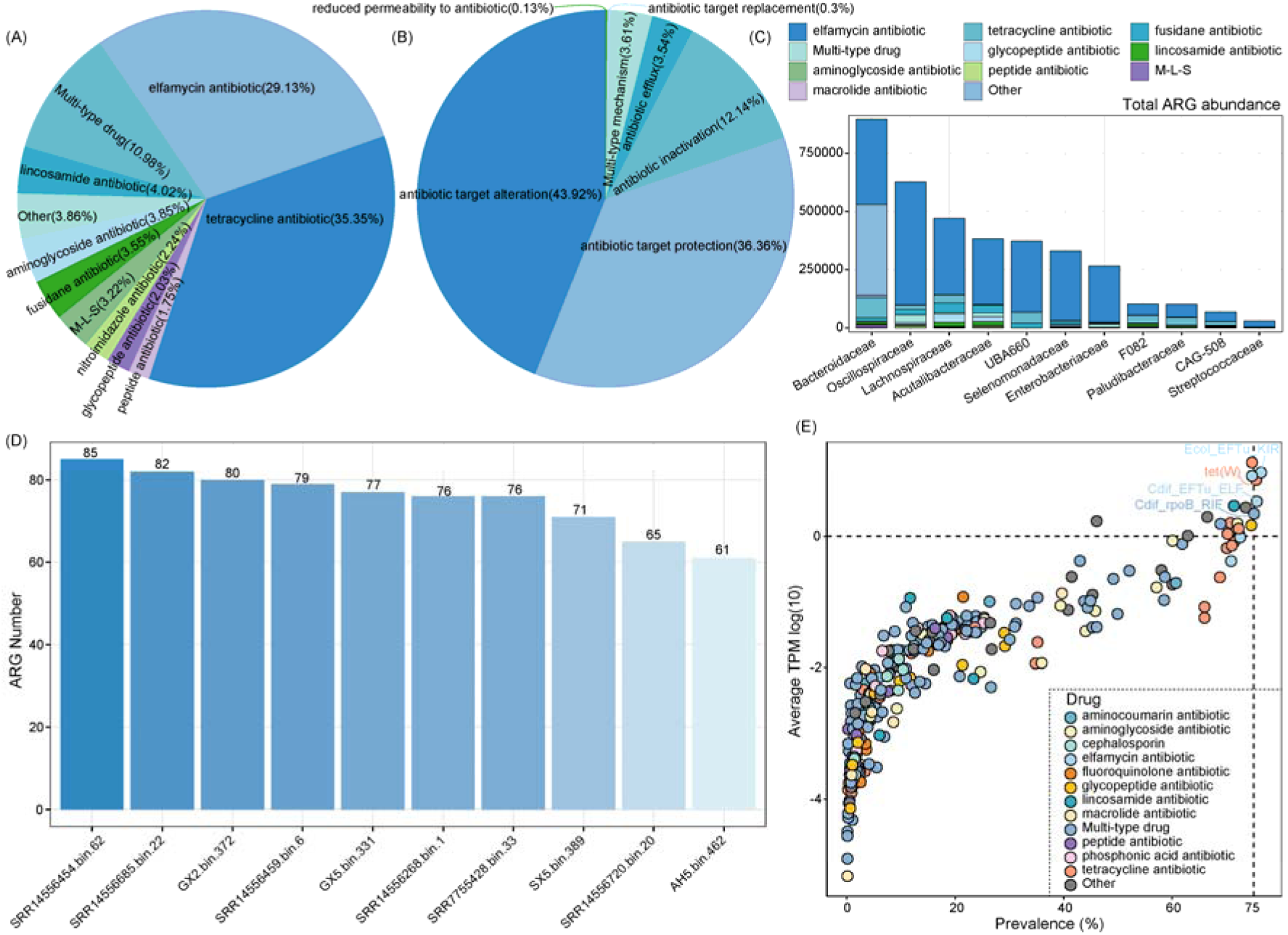
Distribution patterns of ARGs in the Caprinae gut microbiota. (A) Relative abundance of ARGs categorized by resistance phenotypes. (B) Relative abundance of ARGs grouped by resistance mechanisms. (C) The ten bacterial families with the highest relative ARG abundance, with stacked colors indicating associated resistance phenotypes. (D) The ten MAGs harboring the greatest number of ARGs. (E) Prevalence distribution of ARGs, where each point represents an ARG colored by its resistance phenotype.

At the family level, ARGs were predominantly associated with members of *Bacteroidaceae*, *Oscillospiraceae*, and *Lachnospiraceae* (Fig. 2C). At the species level, *Escherichia coli* (*E. coli*) emerged as the top ARG carrier, followed by *UBA2804 sp900319705* and *Enterococcus_B faecium* (Supplementary Fig. 1A). Among the ARG-positive MAGs, 2,091 contained a single ARG, while 605 harbored between 2 and 50 ARGs (Supplementary Fig. 1B). Interestingly, 13 MAGs carried extensive resistance profiles with 51–85 ARGs. Three MAGs, SRR14556454.bin.62, SRR14556685.bin.22, and GX2.bin.372, contained 85, 82, and 80 ARGs respectively, and all were taxonomically assigned to *E. coli* (Fig. 2D; Supplementary Table 2).

Analysis of host range revealed that 116 ARGs were confined to a single microbial host, 135 were found in 2–10 hosts, 72 occurred in 11–50 hosts, and 13 ARGs were broadly distributed across more than 50 host species (Supplementary Fig. 1C). In terms of prevalence, four ARGs were highly widespread, each detected in over 75% of the ARG-positive MAGs: *Ecol_EFTu_KIR* (76.38%), *tet(W)* (75.61%), *Cdif_EFTu_ELF* (75.48%), and *Cdif_rpoB_RIF* (75.10%) (Fig. 2E; Supplementary Table 4)

### VFGs in Caprinae gut microbiota and their link to antibiotic resistance

To explore the landscape of virulence within the gut microbiota of Caprinae animals, we screened 17,023 high-quality MAGs against the Virulence Factor Database (VFDB). This analysis revealed a substantial presence of VFGs: a total of 5,401 VFGs were identified, representing 141 distinct virulence factors (VFs) across 13 virulence factor classes (VFCs). Interestingly, 2,694 MAGs (15.83%) contained at least one VFG (Supplementary Table 5), highlighting the widespread potential for pathogenicity in the Caprinae gut ecosystem. At the individual gene level, the most prevalent VF was elongation factor Tu (*EF-Tu*), found in 53.15% of VFG-positive MAGs, followed by *Capsule* (20.18%) and *GroEL* (8.88%) (Fig. 3A). When categorized by VFCs, the majority of VFGs were linked to adherence systems (63.64%), with immune modulation (21.98%) and motility mechanisms (5.47%) also well represented (Fig. 3B). In terms of prevalence across samples, *EF-Tu* (76.38%), *Capsule* (75.35%), and *GroEL* (74.33%) emerged as the most widespread VFs (Fig. 3C; Supplementary Table 6).

**Fig. 3:**
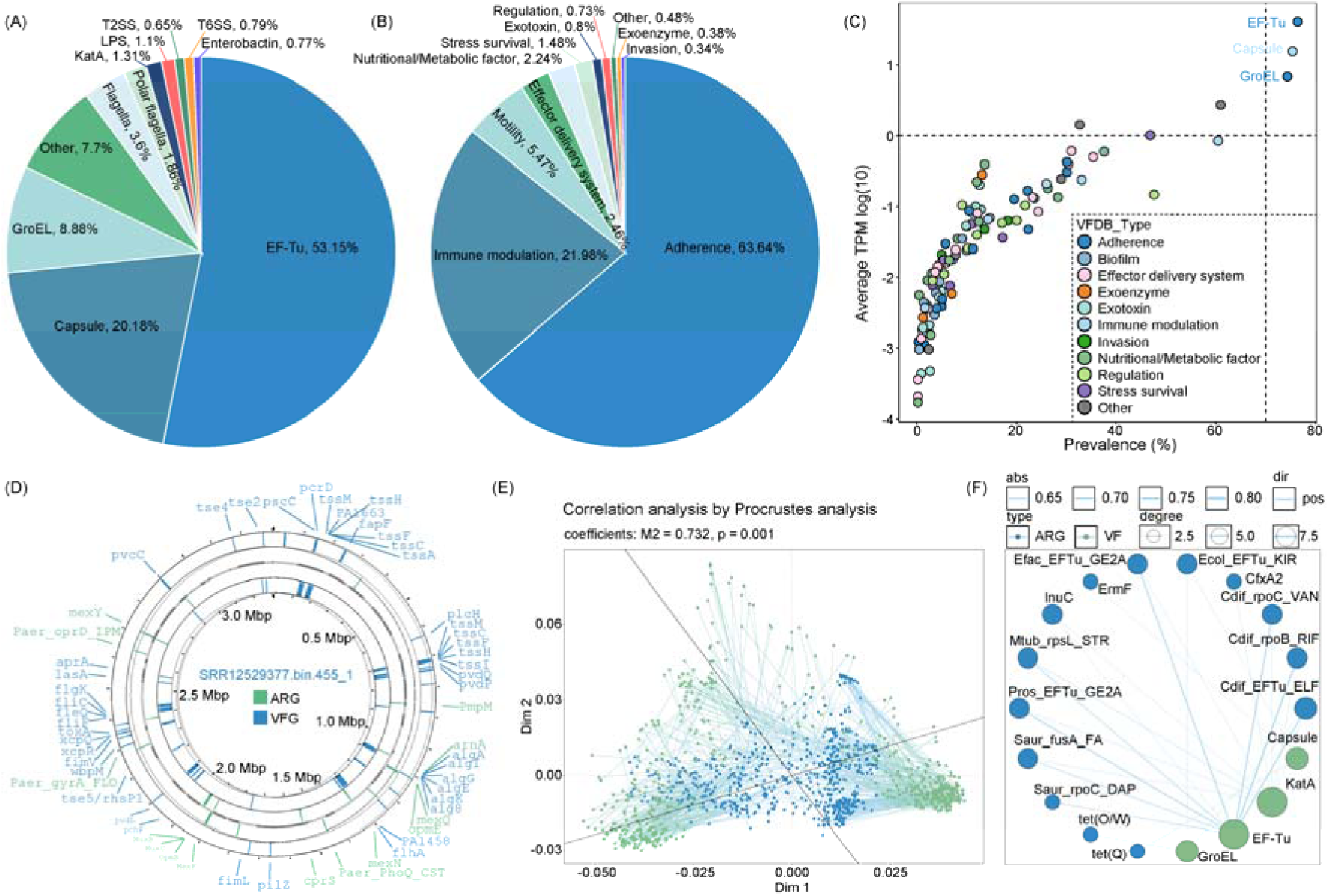
Features of VFs in the Caprinae gut microbiota and their relationship with ARGs. (A) Relative abundance distribution of VFs. (B) Relative abundance of VFCs. (C) Prevalence of VFs, with each dot representing a VF colored by its respective VFC. (D) Circular genome map of SRR12529377.bin.455_1. (E) Procrustes analysis demonstrating a significant correlation between the relative abundances of ARGs and VFGs. (F) Correlation network of the top 20 most prevalent ARGs and VFs, showing significant positive associations (r > 0.6, P < 0.05).

The distribution of VFGs per genome varied significantly. The MAG SRR12529377.bin.455, classified as *Pseudomonas aeruginosa*, harbored the highest number with 199 VFGs, and also carried 31 ARGs—underscoring its pathogenic potential (Fig. 3D). Other highly virulent genomes included *E. coli* strains SRR14556685.bin.22 (107 VFGs) and SRR7755428.bin.33 (104 VFGs) (Supplementary Fig. 2A). Importantly, a strong positive correlation was observed between the abundance of VFGs and ARGs within the microbiota. Diversity analyses revealed significant associations in both Shannon index (R = 0.88, *P* < 2.2 × 10C¹C) and richness index (R = 0.85, *P* < 2.2 × 10C¹C) (Supplementary Fig. 2B, 2C), indicating that communities rich in virulence genes are also rich in resistance genes. Procrustes analysis further reinforced this relationship (M² = 0.732, *P* = 0.001) (Fig. 3E).

Focusing on the 20 most prevalent ARGs and VFs, we identified 24 significant pairwise correlations (r > 0.6, *P* < 0.05) (Supplementary Table 7; Fig. 3F). Interestingly, *GroEL* was strongly associated with eight ARGs, while *KatA* correlated with seven. The strongest single correlation was observed between *GroEL* and *Cdif_rpoB_RIF*, an ARG conferring rifampicin resistance (r = 0.83, padj = 2.83 × 10C¹C²). These findings highlight a potentially synergistic relationship between virulence and antibiotic resistance in the Caprinae gut microbiome, underscoring the importance of integrated surveillance for both traits in animal-associated microbial communities.

### MGEs and phage-mediated resistance in the caprinae gut microbiota

To investigate the potential for horizontal gene transfer and dissemination of antibiotic resistance in Caprinae gut microbiota, we analyzed 17,023 high-quality MAGs against a curated MGE database. This revealed 457 MGE-associated genes across 246 MAGs (1.45%), encompassing 139 unique MGEs grouped into 8 functional types (Supplementary Table 8).

At the gene level, the most abundant MGEs were *2204_tnpA_AB646744.1* (56.23%), *2548_tnpA_U75371.3* (22.35%), and *1897_IS91_MNRK01000014.1* (4.09%) (Fig. 4A). Transposases overwhelmingly dominated the MGE landscape, accounting for 92.87% of all elements, followed by insertion sequences, particularly from the IS91 family (4.80%) (Fig. 4B). In terms of sample prevalence, *2204_tnpA_AB646744.1* was detected in 71.63% of microbiome samples, followed by *2548_tnpA_U75371.3* (50.96%) and *1897_IS91_MNRK01000014.1* (43.13%) (Fig. 4C).

**Fig. 4:**
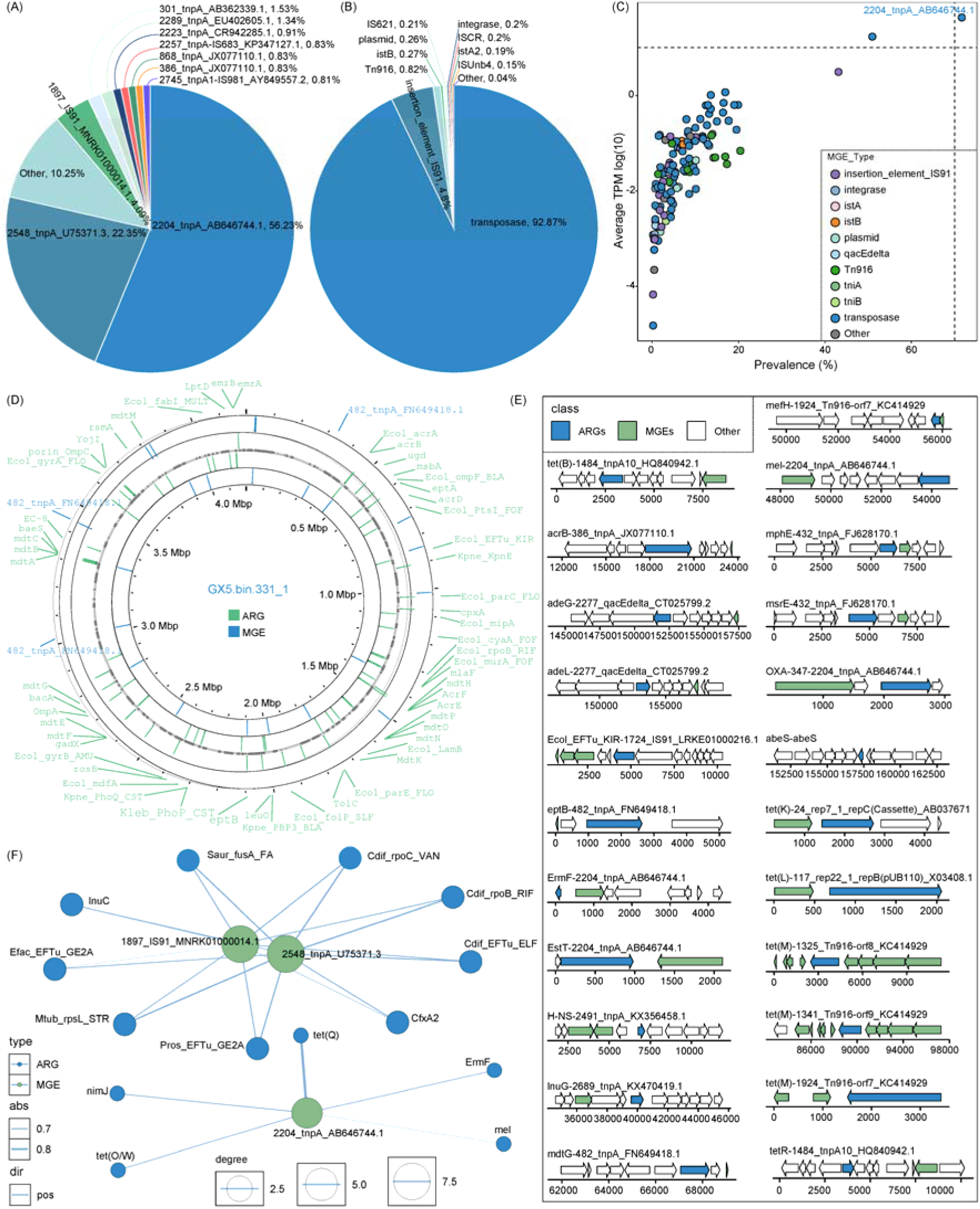
Characteristics of MGEs in the Caprinae gut microbiota and their association with ARGs. (A) Relative abundance of MGEs. (B) Relative abundance distribution across different MGE types. (C) Prevalence distribution of MGEs, with each point representing an MGE colored by its type. (D) Circular genome map of GX5.bin.331_1. (E) Schematic of ARG–MGE co-localization within contigs; arrows indicate gene orientation (right-pointing for forward strand, left-pointing for reverse strand). (F) Correlation network of the top 20 most prevalent ARGs and MGEs, showing significant positive correlations (r > 0.6, P < 0.05).

Some genomes exhibited extensive MGE loads: the *E. coli* genome GX5.bin.331 carried the highest number (22 MGEs), followed by *E. coli* JL4.bin.478, *Dielma fastidiosa* SRR14556658.bin.12, and *E. coli* SX5.bin.389 (each with 13 MGEs) (Supplementary Fig. 3A). Remarkably, GX5.bin.331 also harbored 77 ARGs, underscoring a strong potential for multidrug resistance and genetic mobility (Fig. 4D).

To assess the potential for horizontal gene transfer, we mapped ARGs located within ±5 kb of MGE sequences, uncovering 23 unique MGE-ARG co-localizations. Marked combinations included *tet(B)-1484_tnpA10_HQ840942.1*, *acrB-386_tnpA_JX077110.1*, and *Ecol_EFTu_KIR-1724_IS91_LRKE01000216.1* (Fig. 4E), suggesting these resistance genes may be mobilizable.

Diversity-based correlation analyses further reinforced the connection between MGEs and ARGs. Both the Shannon index (R = 0.83, *P* < 2.2 × 10C¹C) and Richness index (R = 0.91, *P* < 2.2 × 10C¹C) showed strong positive associations (Supplementary Fig. 3B, 3C). Procrustes analysis supported this finding, revealing significant compositional overlap between MGE and ARG profiles (M² = 0.636, *P* = 0.001) (Supplementary Fig. 3D). Among the 20 most prevalent MGEs and ARGs, 25 significant correlations (r > 0.6, *P* < 0.05) were observed, particularly involving three MGEs, *1897_IS91_MNRK01000014.1*, *2548_tnpA_U75371.3*, and *2204_tnpA_AB646744.1*, and 14 ARGs (Fig. 4F; Supplementary Table 10).

Beyond MGEs, bacteriophages may serve as vectors for ARG dissemination. We identified three ARGs within viral genomes: *lnuC* (*Myoviridae*), *Ecol_EFTu_KIR* (*Myoviridae*), and *nimJ* (unclassified) (Supplementary Table 11). These genes confer resistance to lincosamides, elfamycins, and nitroimidazoles, mediated through mechanisms such as antibiotic inactivation and antibiotic target alteration. Predicted bacterial hosts comprised unclassified species from the genera *CAG-196* and *Neisseria*, along with *Bacteroides heparinolyticus*.

These findings underscore the complex and multi-layered architecture of antibiotic resistance in Caprinae gut microbiota, involving both mobile genetic elements and viral-mediated gene transfer. This highlights the need for integrated surveillance approaches to better understand and mitigate the spread of AMR in animal-associated microbial ecosystems.

### Unique and shared ARG profiles in the Caprinae gut microbiota

To characterize host-specific antibiotic resistance profiles, we compared the ARG composition of Caprinae gut microbiota with that of humans and pigs. Across all three hosts, multi-drug resistance emerged as the most dominant resistance type. In Caprinae, multi-drug resistance accounted for 45.83% of ARGs, followed by resistance to tetracyclines (7.74%) and peptide antibiotics (6.85%) (Fig. 5A). Similar patterns were observed in humans (46.31% multi-drug, 6.15% aminoglycoside, 6.01% peptide antibiotics) and pigs (46.39% multi-drug, 8.76% peptide antibiotics, 7.47% aminoglycoside) (Fig. 5B–C). In terms of resistance mechanisms, Caprinae microbiota was primarily shaped by antibiotic efflux (34.52%), target alteration (27.08%), and inactivation (19.05%) (Supplementary Fig. 4A). Human-associated resistomes were dominated by antibiotic inactivation (37.43%), followed by efflux (26.78%) and target alteration (20.49%) (Supplementary Fig. 4B), while pigs exhibited a mechanism profile similar to Caprinae (efflux: 34.28%, target alteration: 27.32%, inactivation: 17.27%) (Supplementary Fig. 4C).

**Fig. 5:**
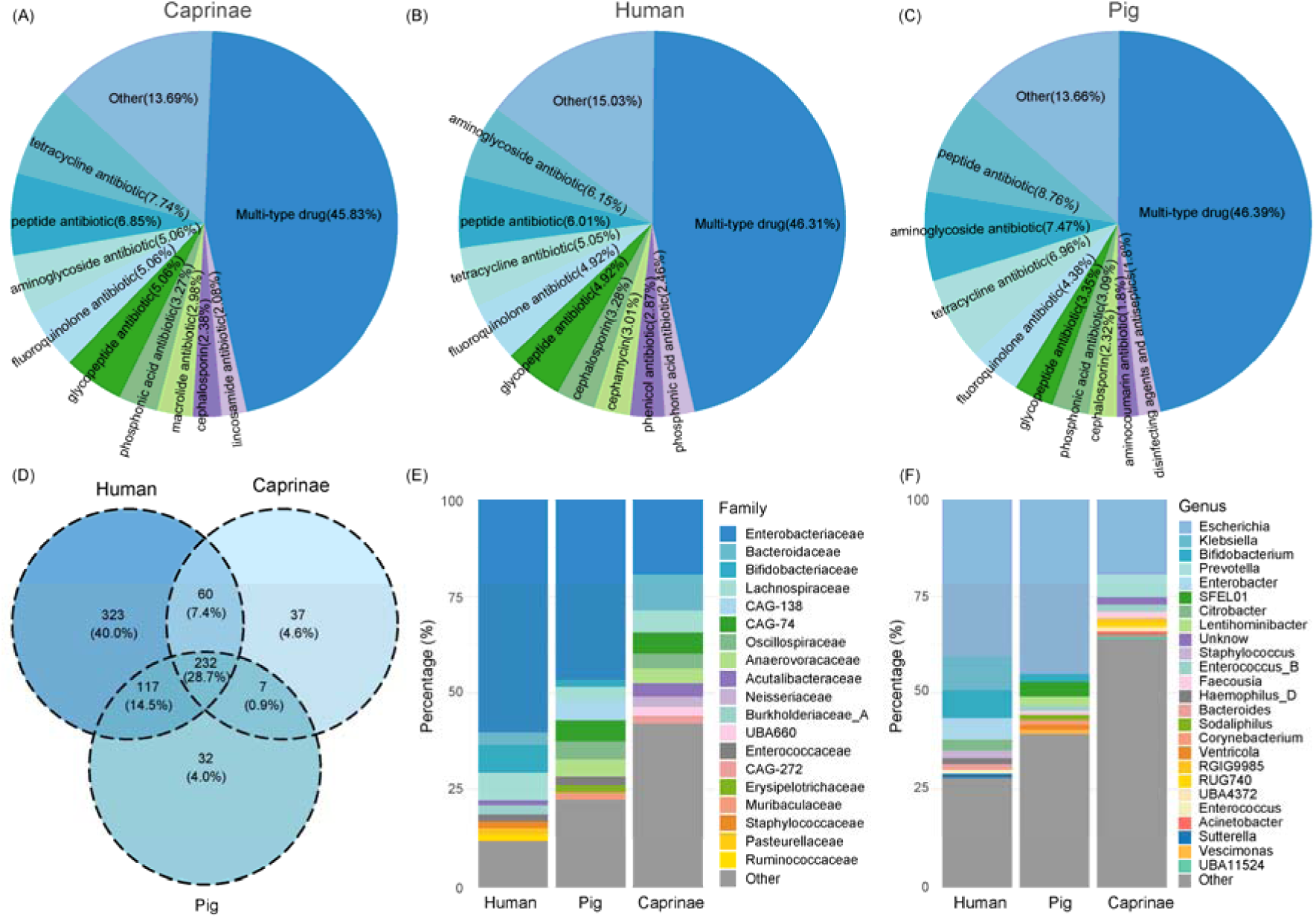
Comparative analysis of ARGs across gut microbiota from Caprinae animals, humans, and pigs. (A) Top ten ARG types with the highest counts in Caprinae gut microbiota. (B) Top ten ARG types predominant in human gut microbiota. (C) Top ten ARG types prevalent in pig gut microbiota. (D) Venn diagram illustrating unique and shared ARGs among Caprinae, pigs, and humans. (E) Taxonomic composition of ARG-carrying microbial genomes at the family level. (F) Taxonomic composition of ARG-carrying microbial genomes at the genus level.

### Host-specific and overlapping ARG profiles

A comparative analysis of ARG repertoires revealed both unique and overlapping resistance genes among hosts (Fig. 5D). Caprinae harbored 37 unique ARGs, in contrast to 323 and 32 unique ARGs in humans and pigs, respectively. Importantly, 4,617 ARGs (292 types) were shared between Caprinae and humans (Supplementary Table 12), and 4,441 ARGs (239 types) were shared between Caprinae and pigs (Supplementary Table 13), underscoring a substantial common resistome across species. At the taxonomic level, Enterobacteriaceae emerged as the primary ARG-carrying family across all hosts (Fig. 5E), with *Escherichia* being the most represented genus (Fig. 5F). However, Caprinae microbiota showed greater diversity in ARG-carrying taxa, exhibiting a broader distribution across genera compared to the more genus-dominated resistomes of humans and pigs.

### Clinically relevant ARGs in caprinae

To assess potential clinical implications, we screened for ARGs conferring resistance to critical antibiotics used against Gram-negative pathogens, including tigecycline, vancomycin, polymyxins, and β-lactams (Supplementary Table 14). Using stringent criteria (identity ≥90%, coverage ≥70%, E-value ≤ 1e−10), BLASTn analysis of the 4,617 ARGs shared between Caprinae and humans identified six high-priority resistance genes: *tetX1*, *tetX4*, *tmexD3*, *vanD*, *vanR*, and *vanS*, resulting in 20 unique matches within Caprinae-derived genomes (Supplementary Table 15). These genes are of concern due to their association with last-line antibiotic resistance and their potential for horizontal transmission.

## Discussion

This study presents a comprehensive metagenomic analysis of ARGs, VFGs, and MGEs in the gut microbiota of Caprinae animals, using 779 publicly available metagenomic datasets. By extending the analysis beyond bacterial genomes to include viral components, particularly bacteriophages, we provide a broader view of the potential vectors involved in ARG dissemination. Our findings highlight Caprinae animals as important reservoirs and potential facilitators of AMR gene persistence and horizontal transfer within host-associated microbial ecosystems.

Among the 17,023 high-quality MAGs, 15.91% harbored at least one ARG, spanning 337 unique resistance phenotypes. The dominant ARG types were associated with tetracycline, elfamycin, and multidrug resistance—a pattern that aligns with the historical use of these antibiotics in livestock production ^4,8^. Notably, several ARGs were detected in over 50 distinct microbial species, suggesting high host plasticity and an increased likelihood of horizontal gene transfer (HGT) ^5,15,16^. Some MAGs carried exceptionally large ARG loads, up to 85 resistance genes in a single genome, with *E. coli* consistently identified as a major ARG carrier. This mirrors trends observed in humans and pigs, reinforcing *E. coli*’s role as a cross-species amplifier of resistance elements and a sentinel for environmental AMR monitoring ^17–19^.

Our analysis identified 5,401 VFGs across 2,694 MAGs, with *EF-Tu*, *Capsule*, and *GroEL* being the most prevalent—corresponding to functions such as adhesion, immune modulation, and motility. Importantly, ARG and VFG profiles showed strong positive correlations in both diversity and abundance, supporting the idea of co-selection under antibiotic pressure ^20^. This linkage between virulence and resistance suggests that antibiotic use may inadvertently select for more pathogenic strains, thereby increasing health risks for both animals and humans ^20,21^. Such dual-functional strains may also disrupt gut microbial equilibrium, potentially reducing resilience to opportunistic pathogens ^22–24^.

Although MGEs were detected in only 1.45% of MAGs, they showed a strong positive correlation with ARG diversity, consistent with their recognized role in mediating HGT. Twenty-three physically linked MGE-ARG pairs were identified within ±5 kb, pointing to a potential for mobilization ^25^. Transposases, particularly *2204_tnpA_AB646744.1*, were highly prevalent, present in over 70% of samples—highlighting their importance as vectors of resistance spread ^26^. In addition, three ARGs were identified in viral genomes, primarily from *Myoviridae* phages, further suggesting that viruses may contribute to ARG dynamics ^27,28^. While current evidence does not yet support a dominant role for phage-mediated ARG transmission, the ecological and evolutionary significance of these findings warrants further experimental investigation ^29,30^.

To assess host specificity and overlap, we compared the Caprinae gut resistome to that of humans and pigs. Caprinae microbiota contained 37 unique ARGs, while 292 and 239 ARG types were shared with humans and pigs, respectively. These overlaps suggest potential routes of genetic exchange, especially in environments such as farms where humans, livestock, and shared water systems intersect ^31,32^. Interestingly, while multidrug resistance dominated across all hosts, Caprinae microbiota primarily relied on efflux pumps and target alteration, contrasting with the enzymatic inactivation mechanisms more prevalent in humans^33^.

Of particular concern, six ARGs shared between Caprinae and human microbiota conferred resistance to clinically important antibiotics such as tigecycline, vancomycin, and polymyxins. The presence of such high-priority resistance genes in animal microbiota raises concerns about livestock serving as environmental reservoirs for ARGs with potential public health implications ^31,34^. While this does not confirm direct transfer to human pathogens, it underscores the importance of continued surveillance under a One Health framework that integrates human, animal, and environmental health.

This study is based on publicly available datasets, which may be influenced by sampling biases in terms of geography, host species, and environmental conditions. Functional annotation relied on existing databases, which may have excluded novel or uncharacterized genes. Furthermore, while sequence co-localization provides insights into potential HGT mechanisms, experimental validation is needed to confirm gene mobility and function. Future research incorporating functional metagenomics, long-read sequencing, and in vitro assays will be essential to fully characterize the mobility and pathogenic potential of key ARGs and MGEs.

## Conclusion

This study offers the first large-scale metagenomic exploration of ARGs, VFGs, and MGEs in the gut microbiota of Caprinae animals. Our findings reveal a broad distribution of ARGs and VFGs across diverse microbial taxa, with *E. coli* emerging as a major resistance reservoir. Significant correlations between ARGs, VFGs, and MGEs suggest co-selection and potential genetic linkage, which may facilitate the persistence and spread of these traits within and between hosts. Importantly, we also detected ARGs in viral genomes, implicating bacteriophages in resistance gene dynamics, although their exact role remains to be fully clarified. Comparative analysis with human and pig gut microbiota revealed substantial overlaps, including ARGs conferring resistance to tigecycline, vancomycin, and polymyxins—antibiotics of critical importance to human health. These findings suggest that Caprinae gut microbiota may act as a hidden reservoir of high-priority ARGs, with implications for both veterinary and human medicine. This work lays a foundation for future studies on the ecological and evolutionary drivers of AMR in livestock and emphasizes the need for integrated AMR monitoring strategies across sectors.

## Methods

### Sample collection

A total of 779 gut metagenomic samples from Caprinae animals were obtained from the NCBI database and the China National Center for Bioinformation (Supplementary Table 1).

### Preprocessing and assembly

The 779 raw metagenomic datasets were initially quality-filtered using Fastp (v0.23.0) ^35^ to remove low-quality reads. Host-derived sequences were then removed by aligning reads against host reference genomes using Bowtie2 (v2.5.0) ^36^. High-quality reads were assembled into contigs with MEGAHIT (v1.2.9) ^37^. The resulting contigs were mapped using BWA (v0.7.17-r1198) ^38^, and sequencing depth was calculated with SAMtools (v1.18) ^39^ alongside the jgi_summarize_BAM_contig_depths script. Metagenome-assembled genomes (MAGs) were binned using MetaBAT2 (v2.15) ^40^ with parameters set to -m 2000, -s 200000, and -- seed 2024. Each bin’s completeness and contamination were assessed using CheckM2 (v1.0.1) ^41^, retaining only those with ≥50% completeness and ≤10% contamination. To eliminate redundancy, high-quality MAGs were dereplicated using dRep (v3.4.3) ^42^ at a 99% average nucleotide identity (ANI) threshold, applying parameters -pa 0.9 and -sa 0.99, resulting in a final non-redundant set of high-quality MAGs.

### Taxonomic annotation and gene prediction of MAGs

Taxonomic classification of the MAGs was conducted using GTDB-Tk (v2.3.2) ^43^ based on the Genome Taxonomy Database (GTDB). Open reading frames (ORFs) were predicted with Prodigal (v2.6.3) ^44^ using default parameters.

### Functional annotation

ARGs were identified by aligning predicted protein sequences against the CARD database ^45^ using DIAMOND (v2.1.8.162) ^46^, applying criteria of at least 80% sequence identity and 80% coverage, with an E-value cutoff of ≤1e-5. Genes conferring resistance to more than two antibiotic classes were classified as multi-drug resistance genes, while those linked to more than two resistance mechanisms were defined as multi-mechanism resistance genes. VFGs were detected by aligning protein sequences against the VFDB ^47^ using the same DIAMOND parameters as for ARGs. Mobile genetic elements (MGEs) were identified by BLASTN (v2.13.0) searches against a customized MGE database curated by Pärnänen et al. ^48^, with alignment thresholds set to an E-value ≤1e-5, minimum 80% identity, and at least 80% query coverage. For estimating the relative abundance of ARGs, VFGs, and MGEs, 20 million clean reads were randomly subsampled from each metagenomic dataset and aligned to the reference gene catalog using Bowtie2 (v2.5.0) ^36^ under default settings. Read counts were normalized to transcripts per million (TPM) for downstream quantitative analyses.

### Identification and processing of viral sequences

To identify viruses potentially involved in the dissemination of AMR, established methods were used to screen contigs >5,000 bp from 2,709 MAGs carrying ARGs ^49,50^. Initially, CheckV (v1.0.1) ^51^ was used to assess the ratio of viral to host genes. Contigs containing more than 10 host genes or where host genes outnumbered viral genes by more than fivefold were excluded. Proviral fragments were also identified using CheckV. Next, multiple viral detection strategies were employed, including (1) viral gene enrichment as determined by CheckV, (2) identification by DeepVirFinder (v1.0.19) ^52^ with a score greater than 0.90 and p-value less than 0.01, and (3) viral classification by VIBRANT (v1.2.1) ^53^ using default parameters. Contigs meeting any of these criteria were retained as putative viral sequences. To remove potential bacterial contamination, BUSCOs ^54^ were used in combination with hmmsearch to detect bacterial single-copy orthologs within the viral candidates. The BUSCO ratio (calculated as the number of BUSCOs divided by total gene count) was computed, and sequences with a ratio of 5% or higher were excluded. The remaining sequences underwent quality assessment with CheckV, and only viral genomes with medium or higher completeness were retained for downstream analyses.

### Viral clustering and representative sequence selection

To eliminate redundancy among viral sequences, viral operational taxonomic units (vOTUs) were constructed following a systematic workflow. First, all viral sequences underwent pairwise alignment using BLASTN with the parameters set to an e-value of 1e-10, a word size of 20, and allowing up to 99,999 alignments. Sequences sharing 95% or greater nucleotide identity over at least 70% of their aligned region were clustered into the same vOTU. Within each vOTU cluster, the longest sequence was chosen as the representative sequence for subsequent analyses.

### Taxonomic clustering of viral populations

To assign viral operational taxonomic units (vOTUs) to genus- and family-level taxonomic groups, pairwise protein sequence comparisons were performed using DIAMOND with an e-value cutoff of 1e-5 and a maximum target sequence count of 99,999. From these alignments, the proportion of shared genes and the average amino acid identity (AAI) between vOTU pairs were calculated. Following the criteria established by Nayfach et al. ^55^, hierarchical clustering was conducted using the Markov Cluster Algorithm (MCL). For family-level clustering, connections with AAI greater than 20% and shared gene content above 10% were retained, applying an inflation parameter of 1.2. For genus-level clustering, more stringent thresholds were used, requiring AAI greater than 50% and shared gene content exceeding 20%, with an inflation parameter of 2.

### Viral taxonomic annotation

Functional annotation of viral genomes was conducted through protein sequence alignments. Predicted proteins were compared against a comprehensive reference database including the Virus-Host DB (May 2024 version), crAss-like viral proteins ^56^, and other published viral protein datasets. Alignments were performed using DIAMOND with parameters set to a minimum identity of 30%, subject coverage of 50%, query coverage of 50%, and a minimum score of 50. For viral genomes with fewer than 30 genes, classification to a viral family required at least 20% of the encoded proteins to match proteins from the same family. For genomes containing 30 or more genes, at least 10 proteins needed to match the same viral family for taxonomic assignment ^49^.

### Correlation analysis

Correlation analyses were conducted using R (v4.4.1). Procrustes analysis was performed with the procrustes function from the ‘vegan’ package. Spearman’s rank correlation coefficient was applied to evaluate the relationships between ARGs and VFGs, as well as between ARGs and MGEs, considering both diversity and abundance profiles.

### Comparative analysis of ARGs in Caprinae, human, and pig gut microbiota

To assess similarities and differences in ARG profiles among Caprinae animals and other key hosts, a comparative analysis was performed using publicly available human and pig gut microbiome datasets. A total of 60,664 human-derived MAGs were obtained from the Integrated Gut Genomes (IGG) database ^57^. Pig-derived metagenomic data were retrieved from the NCBI project PRJNA526405 ^58^ and processed using the same pipeline applied in this study, including quality control, assembly, and genome binning, resulting in 95,445 initial MAGs. All human and pig MAGs underwent consistent downstream processing, including taxonomic classification, gene prediction, quality assessment, and functional annotation. Applying the same quality criteria as for Caprinae MAGs (completeness ≥50% and contamination ≤10%), 36,467 human-derived and 16,394 pig-derived high-quality MAGs were retained. ARG annotation was performed uniformly across all datasets. Using these annotations, we systematically compared ARG distribution patterns across gut microbiota from Caprinae animals, humans, and pigs, with special attention to shared and host-specific ARG phenotypes and their abundance differences.

### Statistical analysis and visualization

Genome structure visualization and target gene annotation were performed using the CGView platform (https://proksee.ca/). Gene arrow diagrams were created with the ‘gggenes’ package (v0.5.1), and network visualizations were generated using the ‘ggraph’ package (v2.1.0). Venn diagrams were plotted with the ‘ggvenn’ package (v0.1.10), while all other figures were produced using the ‘ggplot2’ package (v3.3.6). All statistical analyses were carried out in the R programming environment (v4.4.1).

## Ethical approval

Not applicable.

## Competing interests

The authors declare that there are no financial or personal relationships that could have influenced this study.

## Acknowledgements

X.X.Z. discloses support for the research of this work from the Shandong Province Higher Education Institutions “Youth Innovation Team Plan” [grant number 2022KJ169].

## Data availability

Not applicable.

## Author contributions

**J.W.S.** contributed Writing original draft, Formal analysis, Software, and Visualization; **H.M.E.** contributed Conceptualization and Writing original draft; **L.G.** contributed Supervision and Writing review and editing; **R.L.** contributed Supervision and Writing review and editing; **K.M.S.** contributed Supervision, Formal analysis, Software, and Writing review and editing; **H.L.Y.** contributed Writing original draft, Formal analysis, Software, and Visualization; **H.M.** contributed Writing review and editing; **H.B.N.** contributed Supervision and Writing review and editing; **B.N.C.** contributed Conceptualization, Validation, and Writing review and editing; **X.X.Z.** contributed Conceptualization, Funding acquisition, Validation, Supervision, Software, and Writing review and editing; **X.Y.** contributed Conceptualization, Supervision, and Writing review and editing.

## Supplementary Information

**Additional file 1:**

**Supplementary Figure 1: Host distribution characteristics of ARGs in the gut microbiota of Caprinae animals.** (A) The top ten bacterial species exhibiting the highest relative abundance of ARGs, with stacked colors representing the corresponding resistance phenotypes. (B) Distribution of the number of ARGs carried by individual MAGs. (C) Number of distinct microbial host species associated with each ARG.

**Supplementary Figure 2: Analysis of VFG counts per MAG and their correlation with ARG diversity.** (A) The ten MAGs containing the greatest number of VFGs. (B) Correlation between Shannon diversity indices of ARGs and VFGs. (C) Correlation between richness indices of ARGs and VFGs.

**Supplementary Figure 3: Distribution of MGE-associated gene counts per MAG and correlation with ARG diversity indices.** (A) The ten MAGs harboring the highest number of MGE-associated genes. (B) Correlation between Shannon diversity indices of ARGs and MGE-associated genes. (C) Correlation between richness indices of ARGs and MGE-associated genes.

**Supplementary Figure 4: Comparative analysis of resistance mechanisms in the gut microbiota of Caprinae animals, humans, and pigs.** (A) Dominant resistance mechanisms among the ten most abundant ARGs in Caprinae gut microbiota. (B) Resistance mechanisms associated with the ten most prevalent ARGs in the human gut microbiota. (C) Resistance mechanisms linked to the ten most prevalent ARGs in the pig gut microbiota.

**Additional file 2:**

**Supplementary Tables**

**Supplementary Table 1:** Source details for the 779 Caprinae gut microbiome metagenomic samples included in the study.

**Supplementary Table 2:** Metadata and quality metrics for the 17,023 high-quality MAGs.

**Supplementary Table 3:** Comprehensive annotation of 4,685 ARGs identified in the dataset.

**Supplementary Table 4:** Prevalence and distribution information for detected ARGs across samples.

**Supplementary Table 5:** Detailed annotation of 5,401 VFGs found within the microbial communities.

**Supplementary Table 6:** Prevalence data for VFs across the Caprinae gut microbiomes.

**Supplementary Table 7:** Correlation analysis between the 20 most prevalent ARGs and the 20 most prevalent virulence factors.

**Supplementary Table 8:** Annotation details for 5,401 MGEs detected in the dataset.

**Supplementary Table 9:** Prevalence and distribution of MGEs within the Caprinae gut microbiota.

**Supplementary Table 10:** Correlation analysis between the top 20 ARGs and top 20 MGEs, highlighting potential genetic linkages.

**Supplementary Table 11:** Identification and characterization of ARGs carried within viral genomes.

**Supplementary Table 12:** List of ARGs shared between Caprinae and human gut microbiota.

**Supplementary Table 13:** List of ARGs shared between Caprinae and pig gut microbiota.

**Supplementary Table 14:** Detailed information on shared ARGs conferring resistance to clinically critical antibiotics including tigecycline, vancomycin, polymyxins, and β-lactams.

**Supplementary Table 15:** Identification of shared ARGs conferring resistance to tigecycline, vancomycin, polymyxins, and β-lactams specifically between Caprinae and human gut microbiomes.

## Notes

### Competing Interest Statement

The authors have declared no competing interest.

