## Additional file 1 for "Metagenomic analysis of antimicrobial resistance, virulence, and mobile genetic elements in the gut microbiota of Caprinae species"

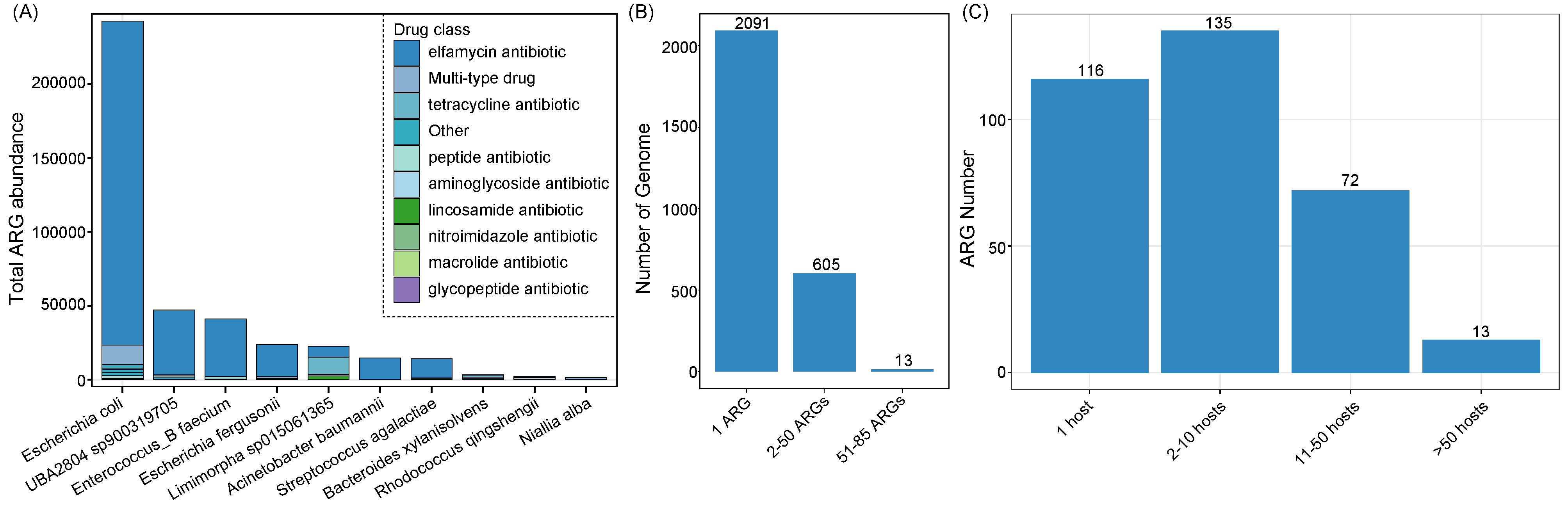

**Supplementary Figure 1:** **Host distribution characteristics of ARGs in the gut microbiota of Caprinae animals.** (A) The top ten bacterial species exhibiting the highest relative abundance of ARGs, with stacked colors representing the corresponding resistance phenotypes. (B) Distribution of the number of ARGs carried by individual MAGs. (C) Number of distinct microbial host species associated with each ARG.

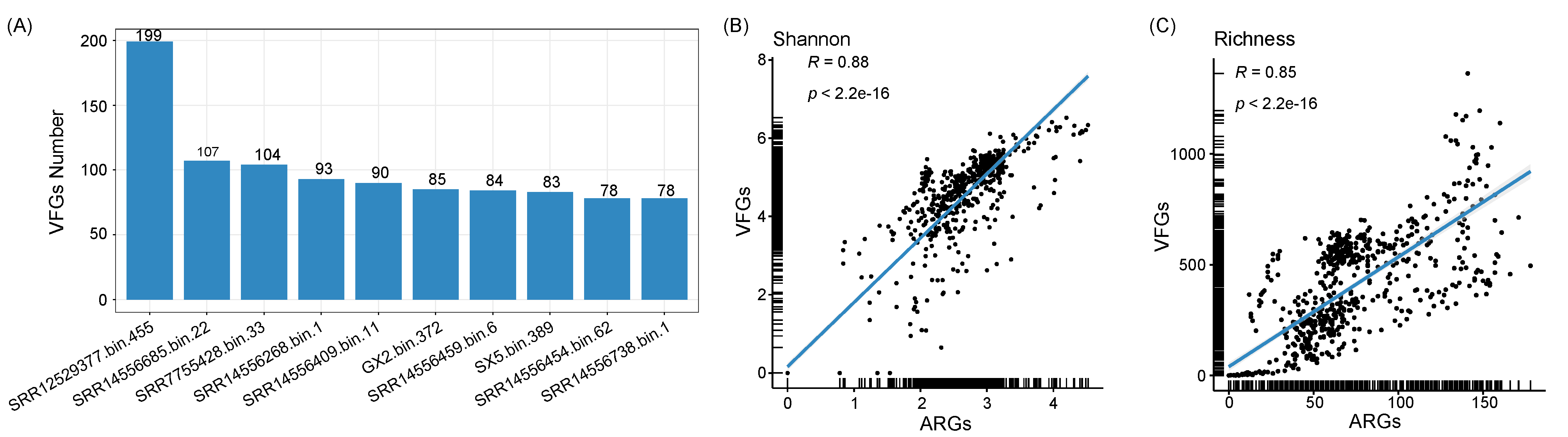

**Supplementary Figure 2:** **Analysis of VFG counts per MAG and their correlation with ARG diversity.** (A) The ten MAGs containing the greatest number of VFGs. (B) Correlation between Shannon diversity indices of ARGs and VFGs. (C) Correlation between richness indices of ARGs and VFGs.

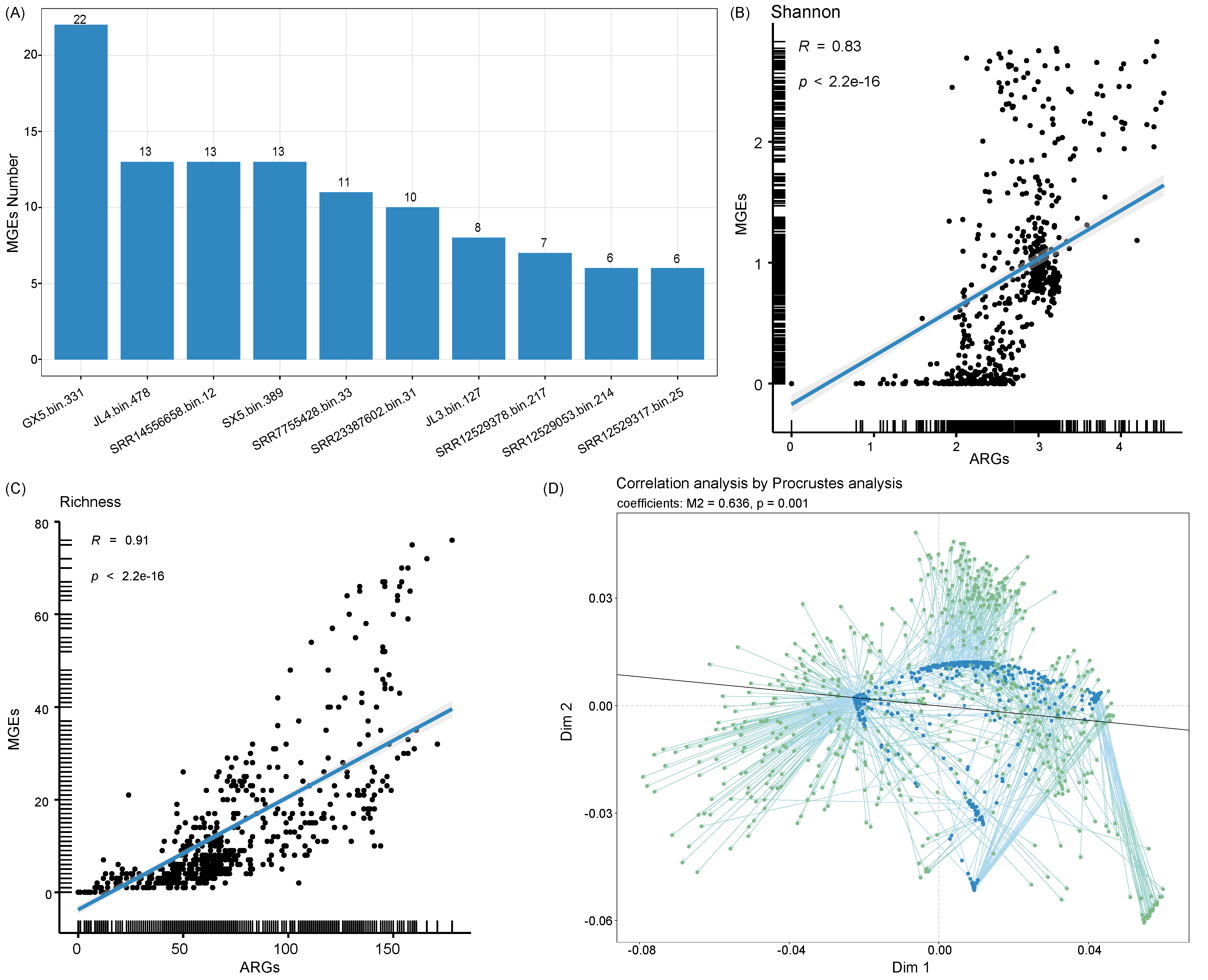

**Supplementary Figure 3:** **Distribution of MGE-associated gene counts per MAG and correlation with ARG diversity indices.** (A) The ten MAGs harboring the highest number of MGE-associated genes. (B) Correlation between Shannon diversity indices of ARGs and MGE-associated genes. (C) Correlation between richness indices of ARGs and MGE-associated genes.

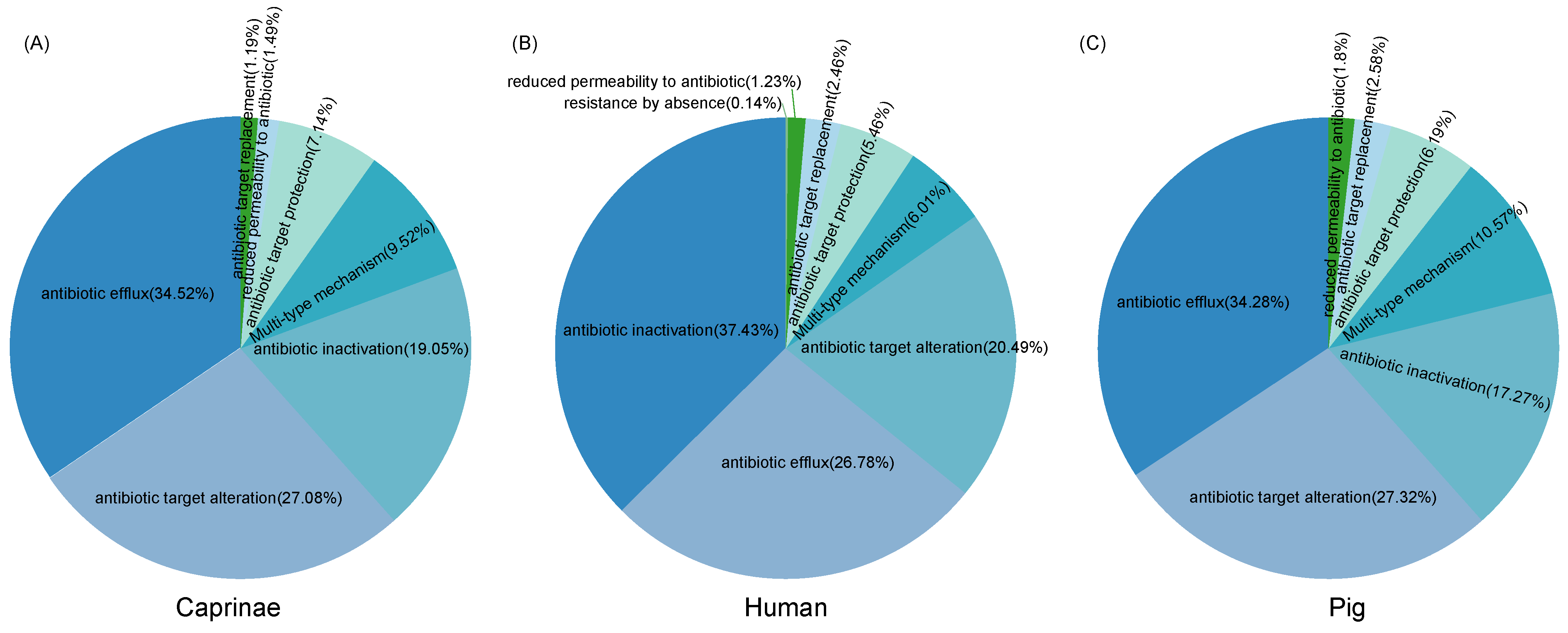

**Supplementary Figure 4:** **Comparative analysis of resistance mechanisms in the gut microbiota of Caprinae animals, humans, and pigs.** (A) Dominant resistance mechanisms among the ten most abundant ARGs in Caprinae gut microbiota. (B) Resistance mechanisms associated with the ten most prevalent ARGs in the human gut microbiota. (C) Resistance mechanisms linked to the ten most prevalent ARGs in the pig gut microbiota.
